# Robotic Imaging and Machine Learning Analysis of Seed Germination: Dissecting the Influence of ABA and DOG1 on Germination Uniformity

**DOI:** 10.1101/2024.05.10.593629

**Authors:** James Eckhardt, Zenan Xing, Vish Subramanian, Aditya Vaidya, Sean Cutler

## Abstract

Seed germination research has evolved over the years, increasingly incorporating technology. Recent advances in phenotyping platforms have increased the accessibility of high throughput phenotyping technologies to more labs, leading to valuable insights into germination biology. These platforms benefit researchers by limiting manual labor and increasing the temporal resolution of imaging. Each of the platforms developed presents unique benefits and challenges, from scalability to price to computing resources. Performing experiments involving thousands of seeds remains a daunting task due to the limitations of current phenotyping platforms and image analysis pipelines. To overcome these challenges, we introduce SPENCER (Seed Phenotype Evaluation and Germination Curve Estimation Robot), a high-throughput phenotyping platform. SPENCER accommodates 32 rectangular petri plates, capable of assessing up to 8000 Arabidopsis seeds per experiment. Our design allows for high quality images while maintaining optimal humidity, crucial for precise germination assessment over longer experiments. The image analysis workflow incorporates advanced image analysis using semantic segmentation models trained for Arabidopsis and lettuce, providing researchers with accessible, reproducible, and efficient tools. We applied SPENCER to investigate the relative roles of DELAY OF GERMINATION 1 (DOG1) and abscisic acid (ABA) in Arabidopsis dormancy. DOG1 mutants exhibited rapid germination, whereas ANT application had a greater impact on the slower-germinating Ler ecotype. Our findings suggest that DOG1 plays a significant role in dormancy, particularly in non-dormant accessions, while ABA’s influence is more pronounced under stress conditions. Additionally, we explored germination uniformity, another agriculurally relevant trait, observing parallels with germination timing. SPENCER offers a powerful and accessible tool for dissecting complex biological traits in conjunction with chemical and genetic manipulations. Its scalability and versatility make it suitable for large-scale genetic and chemical germination screens.

## Introduction

Rising global food demand necessitates enhanced crop productivity, prompting the expansion of agriculture into more challenging environments. Consequently, improving all aspects of growth, development and stress response is essential. The direct impact of seed germination traits on final yield makes them a prime target for enhancement (TeKrony and Egli 1991; W. E. Finch-Savage and Bassel 2016). Stressors such as salt, high temperature and others can negatively affect time to germination and germination uniformity (Daszkowska-Golec 2011). Since germination is important for productivity, experimental approaches and methods are needed to accommodate research. These methods have mainly focused on Arabidopsis thaliana, a model for which there are numerous genetic and genomic, proteomic, metabolomic and transcriptomic resources. Experiments in Arabidopsis have yielded numerous biological findings related to seed traits. The underlying aspects of primary and secondary dormancy, thermoinhibition, stress response and other responses have been well cataloged (Née, Xiang, and Soppe 2017; Huo and Bradford 2015; Finkelstein et al. 2008; William E. Finch-Savage and Leubner-Metzger 2006). Experimental approaches have become more complex. Screens for breeding, genome wide association studies and mutant screening involve thousands of seeds, and classically, germination is counted manually, a time consuming and error prone process. As such, advancements in both software and hardware enabling new high throughput phenotyping methods have been developed to cope with the larger numbers of seeds and the impossibility of the manual counting and manual imaging needed to identify some phenotypic variation underlying all the studied traits (Merieux et al. 2021; Lube et al. 2022; Ligterink and Hilhorst 2017; Colmer et al. 2020).

In seeds, especially Arabidopsis, a plant with small seeds, high throughput phenotyping methods are necessary. Recently, a number of methods have been developed to phenotype both germination and early seedling growth with varying degrees of throughput and temporal resolution. In an early approach, the Busch lab developed a scanner based hardware system coupled with thresholding based software to phenotype root growth (Satbhai, Göschl, and Busch 2017; Slovak et al. 2014). More recently, Merieux et al., 2021 developed hardware and software to quantify germination phenotypes in 96 well plates, graph curves and extract germination kinetics. Image processing was completed with a machine learning approach (Merieux et al. 2021). Lube et al. 2022 also used similar machine learning approaches to extract phenotypic information from images taken from a custom 3D printed high throughput robot (Lube et al. 2022). Modern resources for computer vision, small scale design and manufacturing have all played a part into making high throughput phenotyping more accessible to a wider audience, however we are still in the early stages of this process and there is not a perfect solution to all problems. While compact, a 96 well plate based approach can only accommodate the smallest seeds. The 3D printed approach, while easily accessible, struggles with condensation problems. A scanner based setup lacks the ability to light seeds evenly during growth. More designs are necessary to accelerate high throughput phenotyping. Herein we describe a new **S**eed **P**henotype **E**valuation and Germinatio**n C**urve **E**stimation **R**obot (SPENCER) which deals with some of the previous design challenges and presents a modular modern approach to computer vision and germination kinetics phenotyping. We designed and built a custom robot to accommodate 32 rectangular petri plates, both single well and 6, 8, 12 and 24 well. At maximum capacity for germination assays in Arabidopsis, the robot can phenotype up to 8000 seeds in an experiment. Our approach allows hourly imaging of the plates, and a movable lid ensures high quality images but that the media does not dry even in experiments of up to seven days. To account for the large amount of data generated, we trained two semantic segmentation models in TensorFlow, one for Arabidopsis and one for lettuce, which can be executed and modified in Google Colab. Tensorflow was chosen as one of the most supported and most documentation packages for developing machine learning models (Abadi et al. 2016). Finally, we developed a pipeline in R to accurately extract time to germination data and uniformity data in ways that correctly handle the uncertainty that exists in time to event data. To aid in modularity we did not develop a software package, but rather have made the code available for modification and use. With the use of this system, we quantified the relative contribution of ABA and DELAY OF GERMINATION 1 (DOG1) on time to germination and germination uniformity over a period of after ripening. This illustrates the use of our design and models to aid in answering biological questions about germination kinetics.

## Methods

### Plant material

Arabidopsis lines used in this study were a kind gift from Guillaume Née. The *dog1-2* mutant is in the Col background (Nakabayashi et al. 2012). The NIL DOG1 is a near isogenic line containing the DOG1 allele from Cvi in a Ler background. The *dog1-1* mutant is a dog mutant in the NIL DOG1 background (Bentsink et al. 2006). The genotypes of mutants were confirmed with PCR. Seeds were obtained from plants grown in soil in a growth chamber with a 16/8 light/dark cycle with temperatures of 27°C in 123µmol m^2^s^-1^ light and 21°C in dark. Seeds were harvested and immediately used for experiments or after ripened at 9°C in the dark. All seeds for experiments were harvested at the same time.

Seeds of *Lactuca sativa* ‘Salinas’ were grown in a greenhouse in summer 2022 in Riverside, California. Plants were fully dried and seeds were collected and stored at 9°C in dark for one year before experiments were conducted.

### Germination Assays

Lettuce seeds were surface sterilized and Arabidopsis seeds were gas sterilized immediately before plating. Seeds were grown on filter paper (Ahlstrom 8613) placed on 0.7% agar medium containing ½ Murashige-Skoog (MS) and 10µM ANT or mock untreated control (0.1% v/v DMSO). Petri dishes were immediately transferred to SPENCER. Temperatures ranged from 19-21°C with constant light 98µmol m^2^s^-1^. Seeds were imaged hourly for four days.

### Image Capture and Analysis

Both imaging and robotic control were managed by a single Raspberry Pi, capturing images hourly. These images were then transferred to a computer for lens correction using Photoshop. Post-correction, images were utilized to train two separate U-Net models, one for Lettuce and another for Arabidopsis, manually annotated using the Computer Vision Annotation Tool (CVAT). Our Lettuce dataset comprised 300 images with 3,080 seed objects and 1,612 radicle objects, while the Arabidopsis dataset included 1,212 images with 2,927 seed objects and 2,071 radicle objects. These datasets were partitioned into training (70%), validation (20%), and testing (10%) sets.

The U-Net models, equipped with MobileNetV2 backbones, were trained on TensorFlow with Adam optimizer (learning rate: 1e-4) and Sparse Categorical Crossentropy loss function. Data augmentation, including random horizontal flips, was implemented to bolster model generalization and mitigate overfitting. We optimized training efficiency through prefetching and caching techniques, conducting training in Google Colab to ensure reproducibility and computational efficiency. Model efficacy was gauged using accuracy and loss metrics, and the models were validated on unseen test sets to assess generalization capabilities.

For experimental image processing, images were first aligned using ArUco markers, then cropped to match the annotated image dimensions. The U-Net models were applied to these images for segmentation, followed by stitching the segmented outputs. Seed features were extracted from the initial imaging timepoint, and radicle features were extracted from all timepoints, employing specific algorithms to quantify key morphological attributes. Appendix Figure S7 visually demonstrates the size and distance constraints utilized for seed removal and germination classification.

### Statistical Analysis

ET_50_s (time to 50% germination) were estimated using functions from both *drc* (version 0.1) and *drcte* package (version 1.0.30)(Ritz et al. 2015; Onofri, Mesgaran, and Ritz 2022). The procedure is summarized briefly as follows. Firstly, all possible parametric time-to-event models suggested by previous research were provided as the candidate model list for the time-to-event data from the germination assay(Ritz, Pipper, and Streibig 2013; Onofri, Mesgaran, and Ritz 2022). The models include lognormal, log-logistic, and Weibull I and Weibull II models with two or three parameters. The best-fit model for the data was selected as the one with the lowest AIC (Akaike Information Criterion). After fitting the time-to-event data to the best-fit model, the time to 50% germination (ET_50_) was estimated by the *ED()* function. Pairwise comparisons between different T50s were conducted using the *EDcomp()* function.

## Results

### Robotic Imaging Capabilities

In this study, we developed a novel robotic system, SPENCER (**S**eed **P**henotype **E**valuation and Germinatio**n C**urve **E**stimation **R**obot), which combines a new robotic system for high throughput phenotyping and modern computer vision to accurately estimate germination of multiple species (Fig 1C, S1, S2). This system can capture hourly images of up to thirty-two rectangular petri plates, each of which can accommodate ten rows of Arabidopsis seeds with around twenty-five seeds per row, dispensed by a multichannel pipette. Notably, the versatility of SPENCER extends to other species, as exemplified by its capacity to hold up to 100 lettuce seeds per plate. The design of SPENCER allows for compatibility with different types of rectangular plates, ranging from standard single well plates to eight well rectangular plates, as well as standard 6, 12, and 24 circular well plates. This feature significantly enhances the flexibility in experimental design and broadens the potential use cases. The system incorporates a tightly fitting glass lid, which is robotically removed for optimal image quality every hour thereby mitigating the effects of lid condensation while still maintaining high humidity within the plates and reducing the drying rate. This allows for effective hourly imaging for at least 144 hours. During our standard 96-hour imaging time course, fungal or bacterial growth was not observed on 1/2MS 0.7% agar plates, and rarely problematic even at experiments extending to 144 hours.

**Figure 1.**
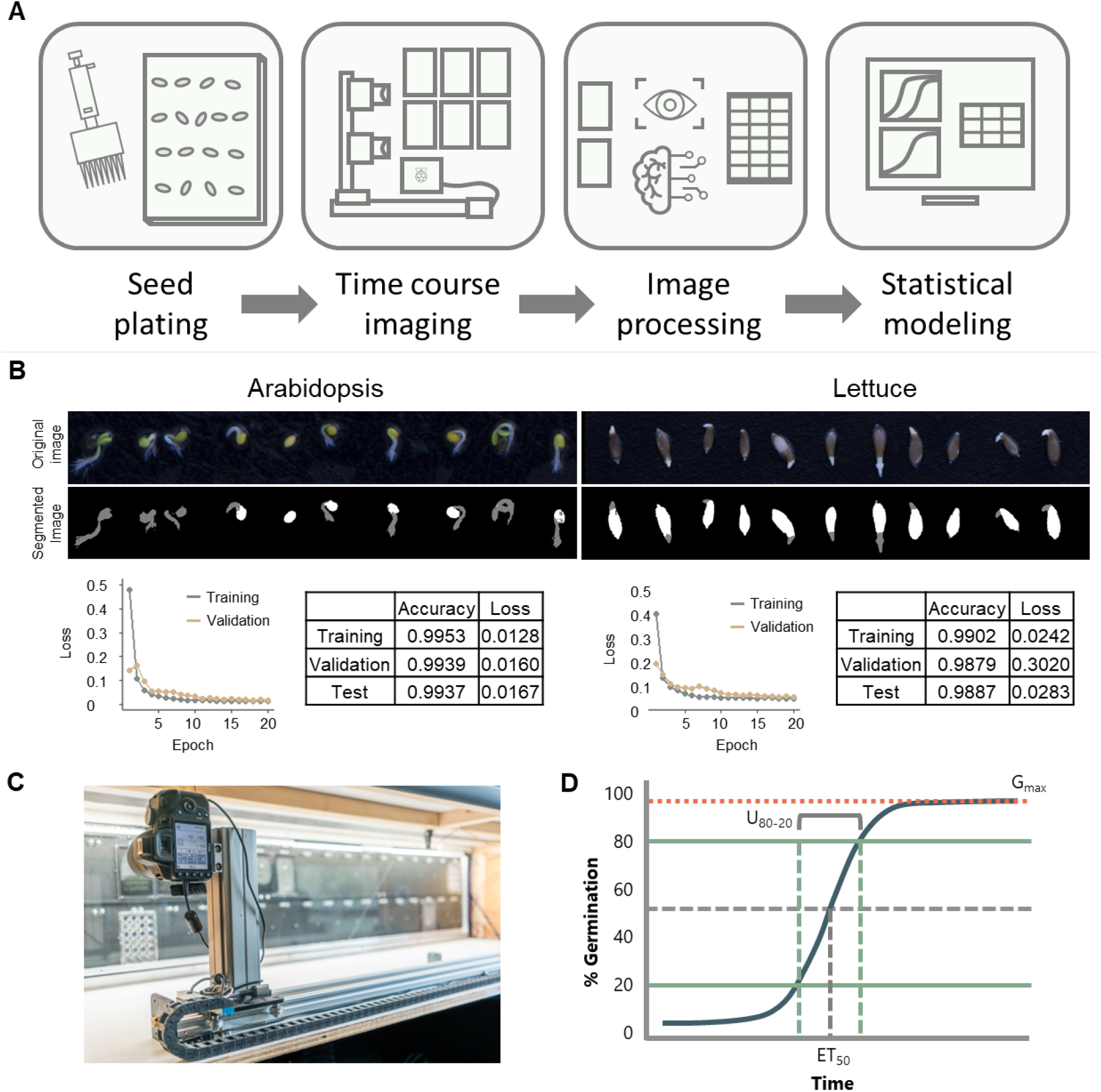
Seed Phenotype EvaluatioN and Curve Estimation Robot (SPENCER). (A) Workflow illustrating steps for imaging, data processing and analysis of seed germination data. Seeds are manually pipetted onto plates, placed in the robot where they are imaged hourly. Images are then segmented with a TensorFlow U-net model. Finally, statistics are estimated in R from the time to event data. (B) Computer vision model performance. Overlays of model outputs for lettuce seeds and Arabidopsis seeds show high model accuracy. This is corroborated by low loss and high accuracy of the models. (C) Image showing SPENCER. (D) Germination curve showing the uniformity (U80-20) and estimated time to 50% germination (ET_50_).

### Efficacy of U-Net Model in Image Segmentation

In this study, we generated two U-Net models, convolutional neural networks, one for Arabidopsis and one for lettuce. We assessed the performance of these models to evaluate their suitability for seed germination semantic segmentation tasks. Both loss and accuracy statistics demonstrate effective model performance. This indicates that the model behaves very similarly to a manual annotation of germination (Fig 1B).

During the training phase, the U-Net model achieved an accuracy of 99.53% for Arabidopsis, indicating its effective learning and adaptation to the training dataset. In the validation phase, the model continued to perform well, achieving an accuracy of 99.39%, suggesting its ability to generalize to new data. In the test phase, the accuracy slightly decreased to 99.37%, which still denotes a reliable performance for unseen data. In lettuce, the model was slightly less accurate and the loss slightly higher than for the Arabidopsis dataset, but still showed the model to be highly effective.

For our loss metrics, we employed the Sparse Categorical Crossentropy loss function, a choice driven by its suitability for multi-class classification with integer labels. This function compares the model’s probabilistic predictions against the actual labels, an efficient method for handling numerous classes. With this function, the model showed a training loss of 0.0128, reflecting a close match between predictions and actual labels. Slight increases in loss were observed during validation (0.0160) and testing (0.0167), a common occurrence when models encounter new data. This indicates that while the model is effectively learning and generalizing, there is a marginal increase in prediction error with new data, which is within expected parameters for this type of model.

### ABA and DOG1 Contributions to Germination Timing and Uniformity

DOG1 is the main contributor to dormancy in Arabidopsis under conditions optimal for germination. Mutants in *dog1* in both Col and Ler backgrounds showed low dormancy even without after-ripening, and ET_50_did not decrease with the amount of after-ripening (Fig. 2, S5). Conversely, the wildtype lines showed significantly slower germination than the dog1 mutants (Appendix table 1). With decreasing ET_50_s as seeds were after-ripened. By two weeks after harvest, Ler and dog1-1 showed only a 1.45 hour ET_50_ difference in the mock treated and a 2 hour ET_50_ difference in the 10µM ANT treated. Col and dog1-2 showed a 06 hour ET_50_ difference in mock treatment and 2.13 hour ET_50_ difference in 10µM ANT treated. This is in contrast to the 12-19 hour ET_50_ differences observed immediately after harvest. This after-ripening effect is consistent with previous literature (Bentsink et al. 2006; Née et al. 2017).

**Figure 2.**
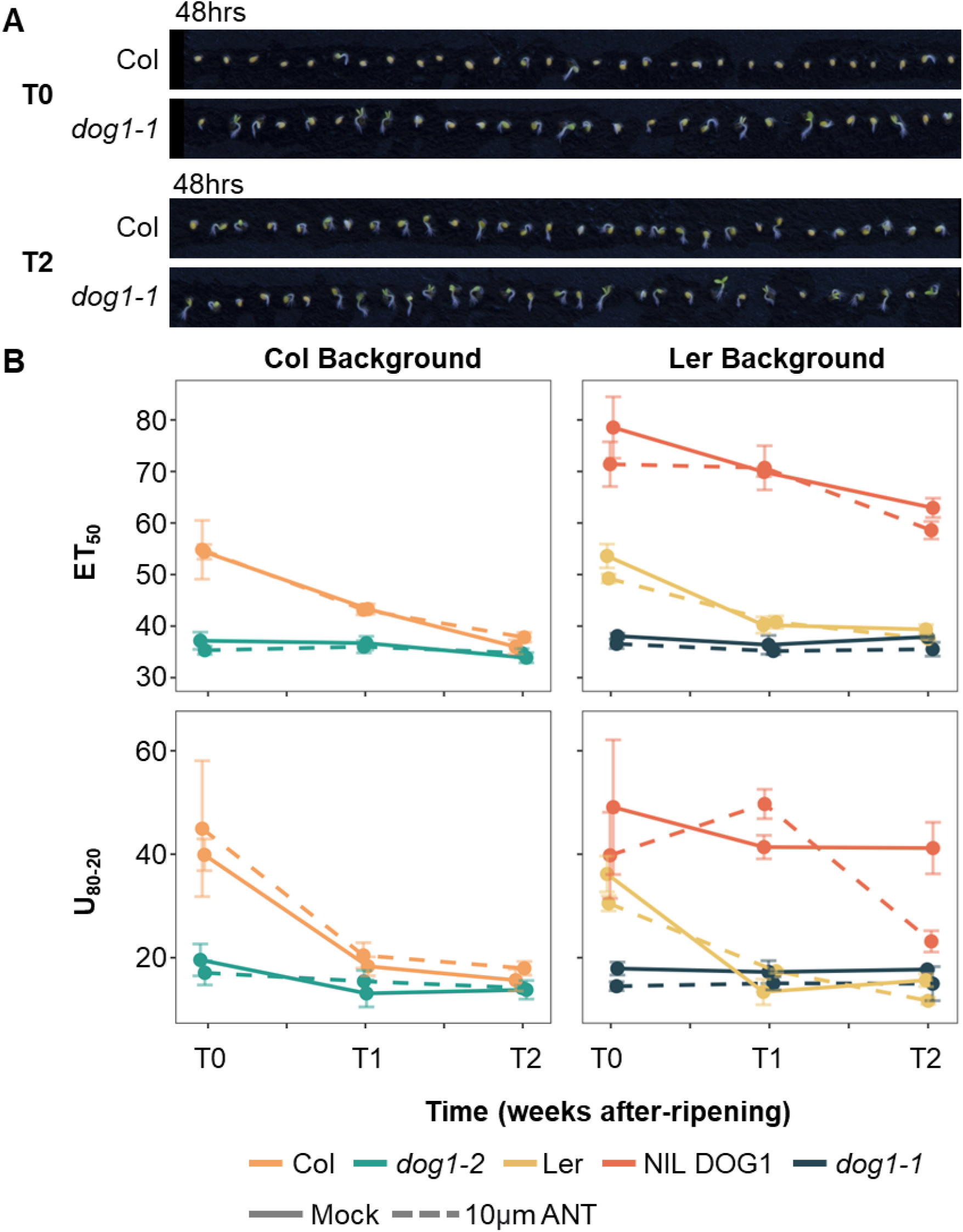
The DOG1 contribution to dormancy decreases during after-ripening. (A) Representative images illustrating the DOG1 contribution to dormancy and after ripening. Images were taken at 48 hours under mock treatment in SPENCER. (B) DOG1 mutants are non dormant, and after-ripening does not alter ET_50_. U80-20 follows similar trends to ET_50_.

While dog1 mutants had an effect on time to germination, blocking ABA signaling did not have a significant impact. Across all lines and times tested, ANT treatment decreased ET_50_ by 1.4hrs (no change in Col backgrounds and 2 hour ET_50_ decrease in Ler backgrounds). Similarly, in lettuce, blocking ABA signaling through ANT application had no effect on time to germination (Fig. S6).

Uniformity followed a very similar trend, albeit with more error, to ET_50_, decreasing as ET_50_ decreased (Fig 2). Data from all treatments and time points were used in a linear regression model to determine the relationship between the two, and ET_50_ was able to explain 85.66% of the variation in uniformity (Multiple R-squared = 0.8566). The association between was highly significant (p <0.001).

## Discussion

Our imaging setup provides a high throughput system to capture high quality images of seeds at a high temporal resolution. During germination, seeds develop quickly, with radicle emergence and growth changing rapidly. As such, a number of labs have developed ways to capture images of seeds and seedlings in the early stages of germination and growth. Each has its unique advantages and disadvantages leading to each being best suited for a different situation. Multiple methods of automatic seed germination scoring and root growth measurements using scanners were developed in multiple labs. These systems can generate high quality images at a high temporal resolution, and can be scaled by increasing the number of scanners. However, precise light control during growth is challenging and the cost of scaling may be prohibitive (Slovak et al. 2014; Teixeira et al. 2007). Teixeira et al. 2007 demonstrated humidity and temperature regulation, but not light control. A more common method of image capture is by camera, and a variety of systems have been developed for this.

The Germinator software required images captured manually under regulated light conditions (Joosen et al. 2010). More recent advances have focused on fully automated image capture. Pavicic et al. 2019 developed a system to automatically capture chlorophyll fluorescence data on a large number (5400) seeds (Pavicic et al. 2019). This method allowed for automated high throughput imaging, however was limited to chlorophyll fluorescence data. SeedGerm provided an inexpensive, scalable solution however struggled with space constraints and image quality issues (Colmer et al. 2020). MultipleXLab features a system of more moderate expense, but reduces some space constraints seen in other systems. It is challenged by uneven lighting and plate condensation issues which need to be manually addressed throughout the course of the experiment (Lube et al. 2022). ScreenSeed is designed to specifically accommodate small seeds such as Arabidopsis, performing experiments in 96 well plates. The obvious advantage of this system is the ability to screen many lines or perform chemical screens. However, the system only accommodates one plate at a time, and only captures reliable data for 24 seeds per well (Merieux et al. 2021). All the methods of image capture, except ScreenSeed, require low throughput seed plating methods to space out seeds for further image analysis, a time consuming process in large scale experiments. SPENCER was designed to reduce some of these limitations, but possesses its own advantages and disadvantages. SPENCER was designed to accommodate a large number of plates (32) in a single experiment allowing for large scale experiments. Unlike the other methods presented, the lighting during growth is evenly distributed to all seeds, and during imaging, the lighting switches to an even diffuse lighting scheme for optimal imaging. Additionally, the lid covering the plates is removed during each imaging step to improve image quality and nullify any issues that condensation would cause. While the time course necessary for the experiments here were only four days, tests show that agar remains hydrated for at least seven days. Together these allow for a fully automated system to capture high quality images without any oversight over the course of large scale experiments. That said, SPENCER has its own set of disadvantages. Firstly, seed plating is still a time consuming and low throughput process. This issue is present in most of the currently published systems, but it continues to be the major bottleneck. High end solutions exist, but some promising accessible methods are being developed. The Araprinter from the Alonso-Stepanova lab (https://diyplantbiology.wordpress.ncsu.edu/) is a cost effective solution, however, plating speed is a limiting factor. There is also some promise using liquid handling robots (OT-2 Robot) to plate seeds. The second issue, although minor, is the possibility of contamination due to removing the glass plate cover during imaging steps. This however, can largely be avoided in experiments not involving sucrose in the media. Finally, SPENCER uses a custom machined aluminum back panel to hold the plates, and although the metal and machining cost can be comparable to other systems described, accessibility to machining may present an issue. All other hardware is easily purchasable and cost effective, making only the back plate a challenge. Other engineering solutions are also possible, but were not considered due to accessibility and ease of machining.

In addition to hardware, code was also developed to process images in a high throughput manner (Fig. S3, S4). We developed a pipeline to process the images, perform semantic segmentation, extract feature information and then model germination and extract kinetics in R (Fig. 1D). Notably, all imaging processing and segmentation steps can be run in Google Colab without the need for computing cores or other hardware. We chose to build our code using languages familiar to biologists, R and Python, with modern packages with readily available resources and support, Tensorflow and OpenCV. The code was built with modularity in mind. While an effective phenotyping robot, it is unlikely SPENCER will be reproduced, however it is much more likely parts from the code will be reproduced. Other phenotyping platforms have chosen different approaches to code accessibility, from software packages to approaches with modularity and improvement in mind. As time has progressed so has the use of machine learning models and approaches over both manual and image segmentation. This is a rapidly growing and changing field, new models are being developed, new best practices are developed and current methods and models will be obsolete. In this rapidly evolving field, a software package is then less helpful to the community than simple modular code. We designed our code to be run in Colab, so that it is accessible to be run by anyone with internet connectivity. Each step of the image processing and analysis pipeline can be taken and repurposed. For example, if a different model is preferred for the user’s use case, this code chunk can be removed and the new code can replace it. Additionally, with the increased accessibility to coding brought about by ChatGPT this can be accomplished more easily by those it was previously inaccessible to. Again, it is important to review some disadvantages of this system. As opposed to a graphical user interface, some knowledge of coding must accompany any use of the Python image analysis code or the R germination modeling code. Additionally, due to resource constraints in Google Colab, the image analysis can take longer than a high performance computing core. Four day experiments take no longer than an overnight processing time, so this is not highly limiting. Finally, if a researcher wanted to run the pipeline on a set of images other than those taken in SPENCER, a new training dataset would be needed to generate another model. This can be a time consuming process based on the complexity of the image dataset, but it is only becoming easier with AI aided image annotation.

We applied this high throughput phenotyping platform and image analysis pipeline to determine the effect of DOG1 and ABA on dormancy in Arabidopsis. DOG1 is a well known dormancy regulating factor, however the molecular function is unsolved. Previous research has shown that mutations in DOG1 completely abolish any seed dormancy (Bentsink et al. 2006; Née et al. 2017). Even when planted immediately after harvest, DOG1 mutants in the Columbia and NIL Cvi background were non dormant. In wildtype plants, the amount of DOG1 protein in seeds determines the amount of after ripening before dormancy release (Nakabayashi et al. 2012). Nakabayashi et al. 2012 propose this happens in a process largely independent of ABA. However ABA is also highly important in dormancy and has been intensively studied. ABA is perceived by the PYR/PYL protein receptors, which upon ABA binding, bind to and inhibit protein phosphatase 2C (PP2C) clad A proteins (Park et al. 2009; Fujii et al. 2009). These proteins then are unable to inhibit the activity of SNF1-related protein kinase 2 (SnRK2) proteins, which show reduced dormancy in mutants (Nakashima et al. 2009). Antabactin, a potent ABA receptor antagonist, restores germination of seeds that have their germination inhibited by ABA. When applied, it reduces ABA signaling (Vaidya et al. 2021). This chemical, when applied in coordination with DOG1 mutants, can allow us to understand seed germination kinetics in the absence of ABA and DOG1 action and identify the relative importance of each factor in determining dormancy, uniformity and time to germination. We observed the germination kinetics of five lines, Col, *dog1-2*, Ler, NIL DOG1, *dog1-1*, over a period of three weeks. As seen in previous literature, the *dog1-1* and *dog1-2* mutants show rapid germination and little change as they are after ripened. Both Col, Ler and NIL DOG1 show reduced dormancy over time, as previously shown (Bentsink et al. 2006; Née et al. 2017), with NIL DOG1 being the most dormant. Col and *dog1-2* did not show any significant change in time to germination or uniformity when ABA signaling was blocked through ANT application, but as expected, Ler was the slower germinating ecotype and showed more ABA effect, albeit not significant. This shows that in generally non dormant accessions, DOG1 plays a major role in after ripening and dormancy, whereas ABA plays only a minor role. ABA may be much more important in regulating dormancy response under stress responses. As seen in thermoinhibition, application of ANT in multiple species has significantly decreased time to germination (Vaidya et al. 2021). This work illustrates the relative importance of DOG1 and ABA in dormancy and after ripening.

Little is known about uniformity and the relative importance of these two factors in regulating uniformity. Germination uniformity, which has been proposed as a bet hedging strategy, was measured by SPENCER. It exhibits parallel patterns to the timing of germination, and regression analysis showed that it was significantly related with most variation in uniformity being explained by ET_50_.

Taken together, this work shows the use of a high throughput phenotyping and image analysis robot in teasing apart complex biological traits in conjunction with chemical and genetic manipulations. The success of this approach opens new avenues for future research, especially in the field of functional genomics and in conjunction with quantitative genomic methods. The integration of high-throughput imaging with detailed genetic and chemical analyses allows for a more nuanced understanding of the underlying mechanisms controlling seed germination and dormancy. The robot or experimental procedures can be modified to allow for modifications of lighting and chemical content allowing for more complex biological questions to be asked. Additionally, the scalability of this approach makes it suitable for large-scale genetic screens on mutant populations or panels of natural variation, enabling the identification of novel genes and pathways involved in plant development and stress responses.

Further advancements in image processing algorithms, particularly those incorporating machine learning and AI, could enhance the accuracy and efficiency of data analysis. This would allow for the extraction of more complex phenotypic data, leading to a deeper understanding of plant biology. Another promising direction is the integration of this technology with other omics approaches, such as transcriptomics, proteomics, and metabolomics, to provide a holistic view of plant responses at different levels of biological organization.

Moreover, the modularity and accessibility of the system’s software make it a valuable tool for educational and research institutions with limited resources. By enabling more researchers to engage in high-throughput phenotyping, this technology democratizes access to cutting-edge research tools, thereby aiding future research.

In summary, this work not only demonstrates the power of combining robotics, imaging, genetics, and chemistry in plant research but also sets the stage for future explorations that could significantly advance our understanding of plant biology and improve agricultural practices.

## Acknowledgements

This work was partially supported by NSF grant 1656890.

