## Supplemental Figures for "Robotic Imaging and Machine Learning Analysis of Seed Germination: Dissecting the Influence of ABA and DOG1 on Germination Uniformity"

### Slide 1
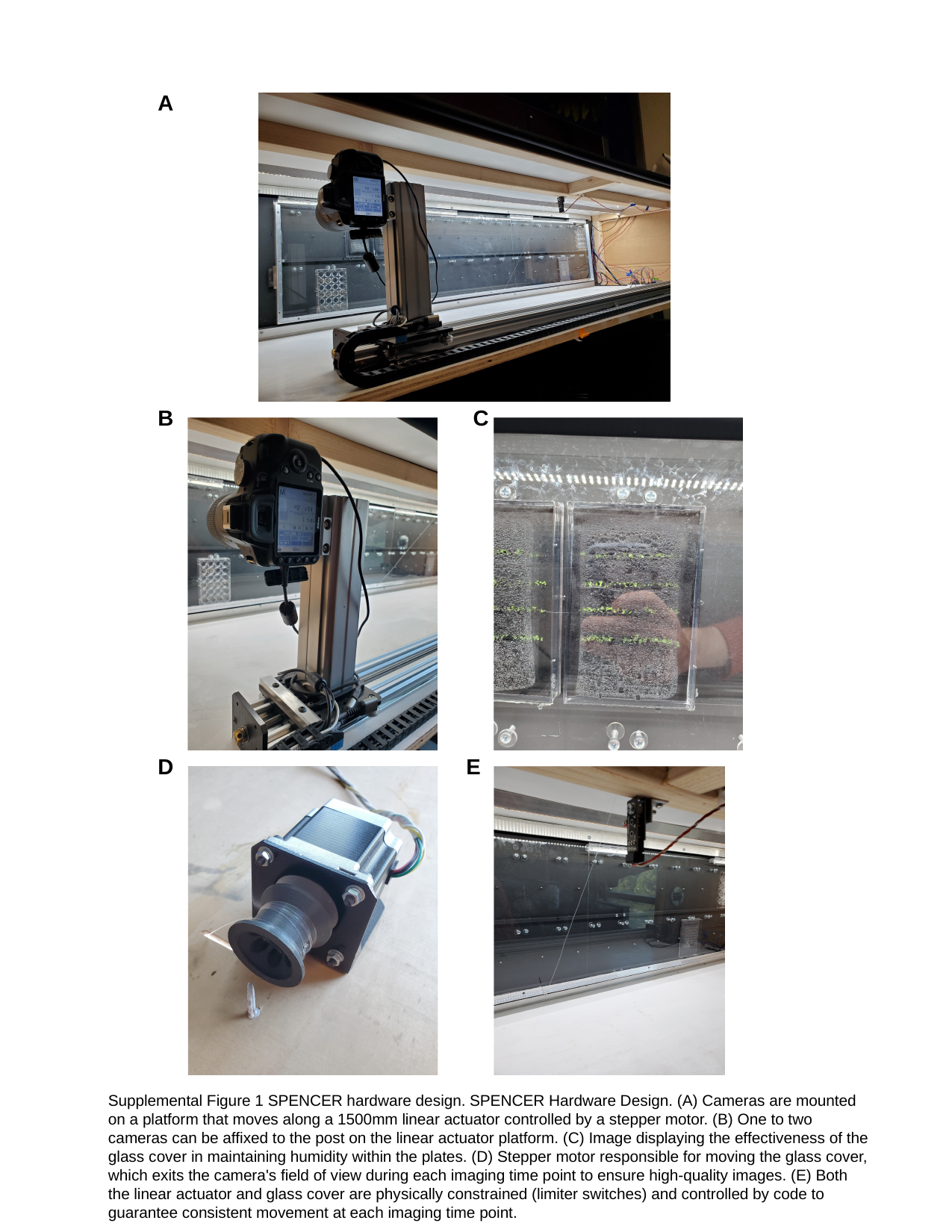

A
B
C
D
E
Supplemental Figure 1 SPENCER hardware design. SPENCER Hardware Design. (A) Cameras are mounted on a platform that moves along a 1500mm linear actuator controlled by a stepper motor. (B) One to two cameras can be affixed to the post on the linear actuator platform. (C) Image displaying the effectiveness of the glass cover in maintaining humidity within the plates. (D) Stepper motor responsible for moving the glass cover, which exits the camera's field of view during each imaging time point to ensure high-quality images. (E) Both the linear actuator and glass cover are physically constrained (limiter switches) and controlled by code to guarantee consistent movement at each imaging time point.

### Slide 2
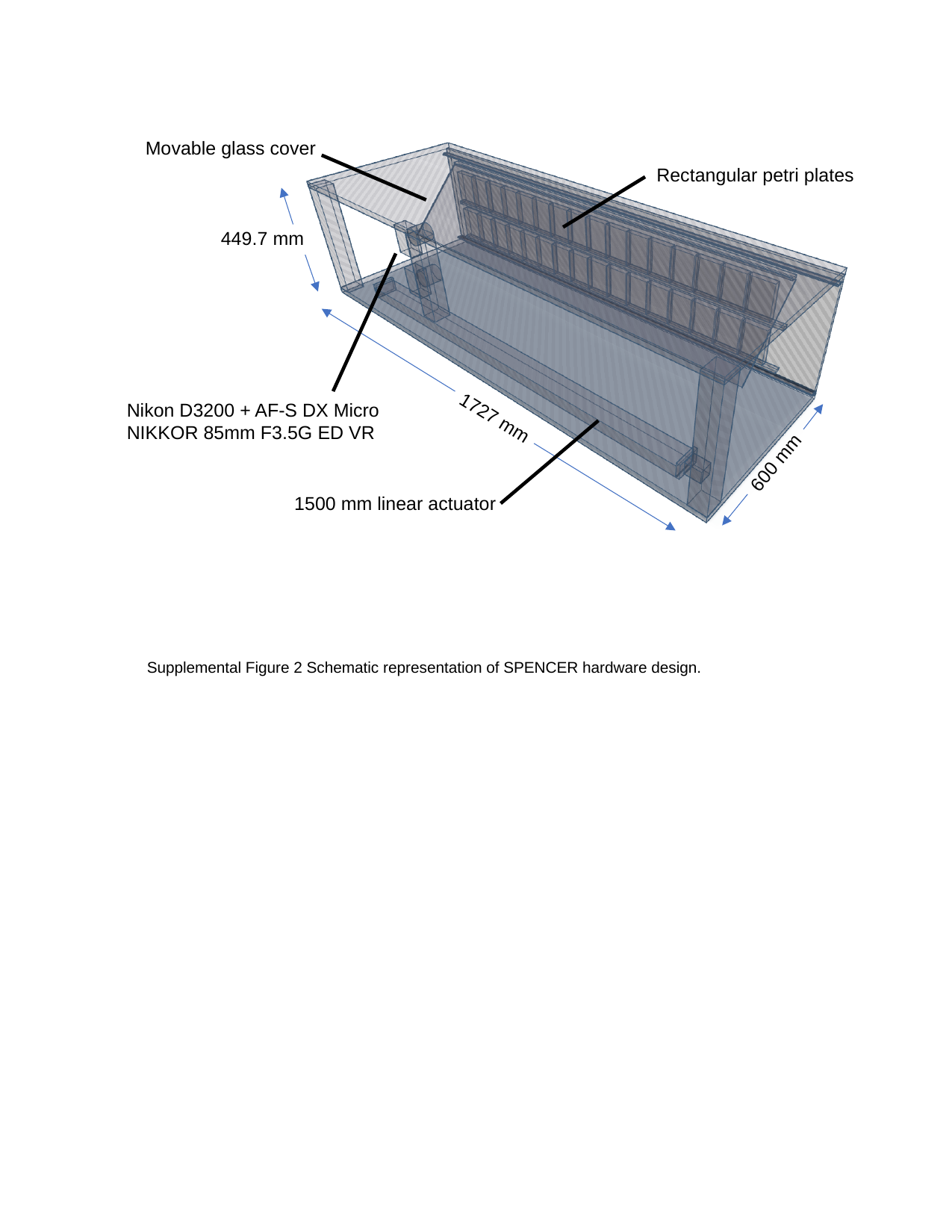

Movable glass cover
Rectangular petri plates
449.7 mm
Nikon D3200 + AF-S DX Micro NIKKOR 85mm F3.5G ED VR
1727 mm
600 mm
1500 mm linear actuator
Supplemental Figure 2 Schematic representation of SPENCER hardware design.

### Slide 3
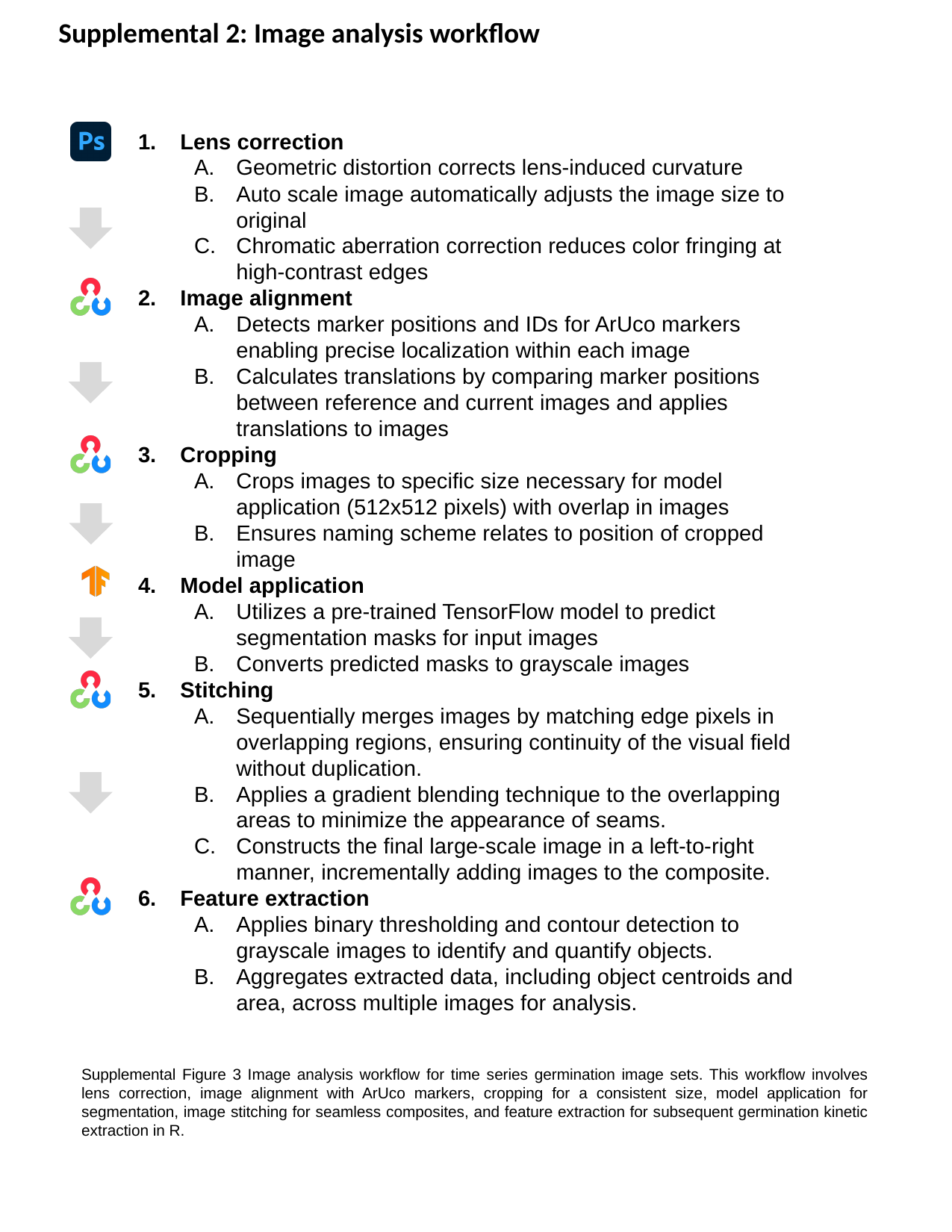

Supplemental 2: Image analysis workflow
Lens correction
Geometric distortion corrects lens-induced curvature
Auto scale image automatically adjusts the image size to original
Chromatic aberration correction reduces color fringing at high-contrast edges
Image alignment
Detects marker positions and IDs for ArUco markers enabling precise localization within each image
Calculates translations by comparing marker positions between reference and current images and applies translations to images
Cropping
Crops images to specific size necessary for model application (512x512 pixels) with overlap in images
Ensures naming scheme relates to position of cropped image
Model application
Utilizes a pre-trained TensorFlow model to predict segmentation masks for input images
Converts predicted masks to grayscale images
Stitching
Sequentially merges images by matching edge pixels in overlapping regions, ensuring continuity of the visual field without duplication.
Applies a gradient blending technique to the overlapping areas to minimize the appearance of seams.
Constructs the final large-scale image in a left-to-right manner, incrementally adding images to the composite.
Feature extraction
Applies binary thresholding and contour detection to grayscale images to identify and quantify objects.
Aggregates extracted data, including object centroids and area, across multiple images for analysis.
Supplemental Figure 3 Image analysis workflow for time series germination image sets. This workflow involves lens correction, image alignment with ArUco markers, cropping for a consistent size, model application for segmentation, image stitching for seamless composites, and feature extraction for subsequent germination kinetic extraction in R.

### Slide 4
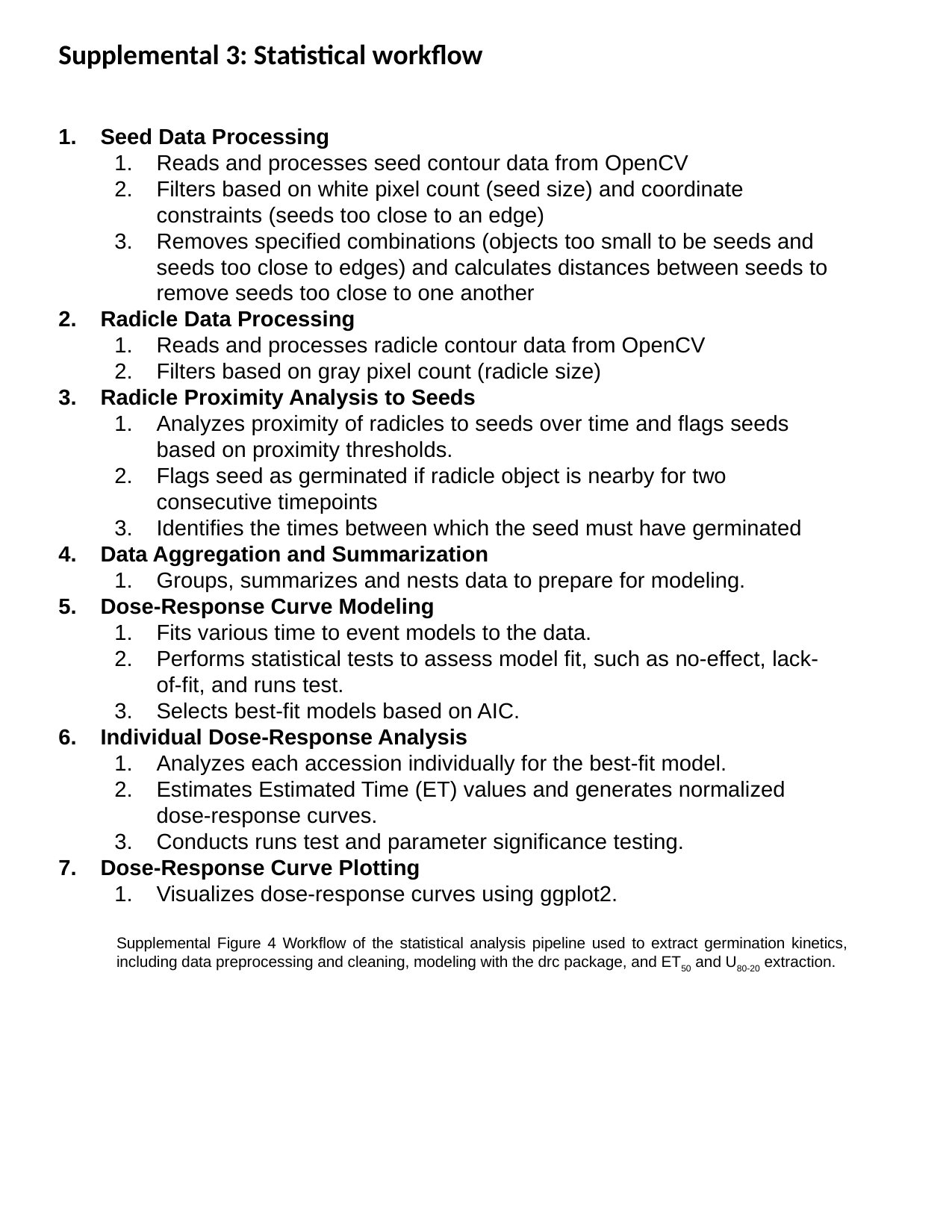

Supplemental 3: Statistical workflow
Seed Data Processing
Reads and processes seed contour data from OpenCV
Filters based on white pixel count (seed size) and coordinate constraints (seeds too close to an edge)
Removes specified combinations (objects too small to be seeds and seeds too close to edges) and calculates distances between seeds to remove seeds too close to one another
Radicle Data Processing
Reads and processes radicle contour data from OpenCV
Filters based on gray pixel count (radicle size)
Radicle Proximity Analysis to Seeds
Analyzes proximity of radicles to seeds over time and flags seeds based on proximity thresholds.
Flags seed as germinated if radicle object is nearby for two consecutive timepoints
Identifies the times between which the seed must have germinated
Data Aggregation and Summarization
Groups, summarizes and nests data to prepare for modeling.
Dose-Response Curve Modeling
Fits various time to event models to the data.
Performs statistical tests to assess model fit, such as no-effect, lack-of-fit, and runs test.
Selects best-fit models based on AIC.
Individual Dose-Response Analysis
Analyzes each accession individually for the best-fit model.
Estimates Estimated Time (ET) values and generates normalized dose-response curves.
Conducts runs test and parameter significance testing.
Dose-Response Curve Plotting
Visualizes dose-response curves using ggplot2.
Supplemental Figure 4 Workflow of the statistical analysis pipeline used to extract germination kinetics, including data preprocessing and cleaning, modeling with the drc package, and ET_50_ and U_80-20_ extraction.

### Slide 5
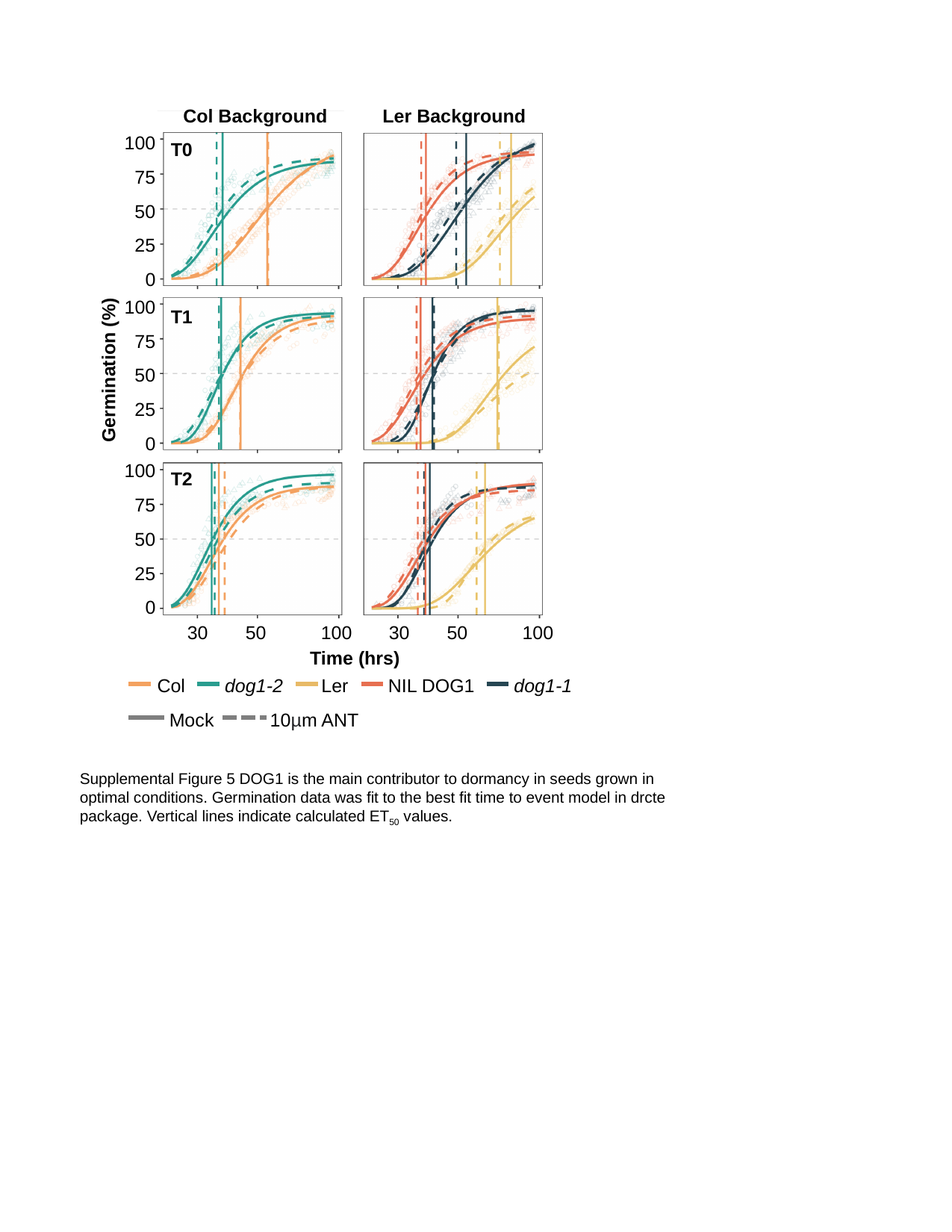

Col Background
Ler Background
100
T0
75
50
25
0
100
T1
75
Germination (%)
50
25
0
100
T2
75
50
25
0
30
50
100
30
50
100
Time (hrs)
Col
dog1-2
Ler
NIL DOG1
dog1-1
Mock
10µm ANT
Supplemental Figure 5 DOG1 is the main contributor to dormancy in seeds grown in optimal conditions. Germination data was fit to the best fit time to event model in drcte package. Vertical lines indicate calculated ET_50_ values.

### Slide 6
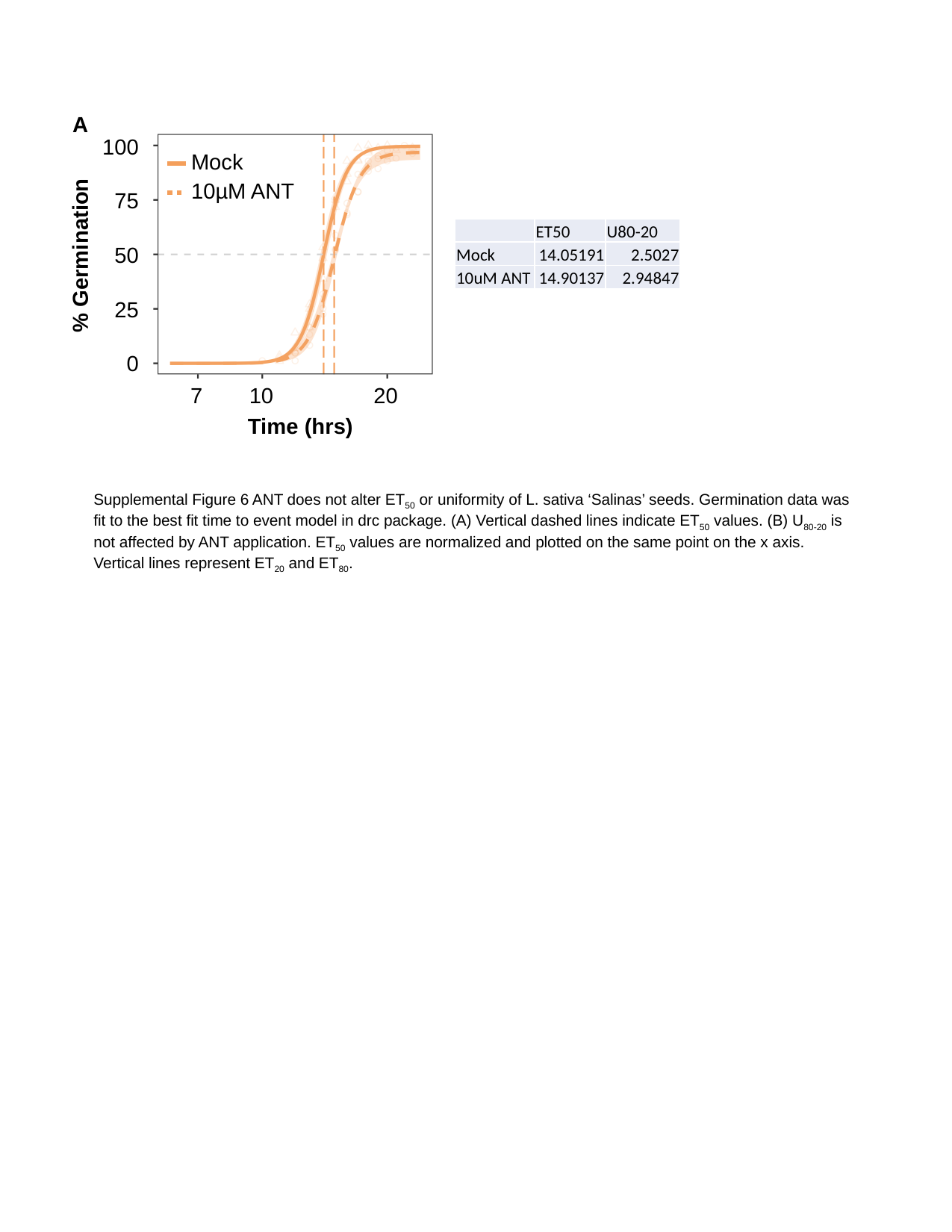

A
100
Mock
10µM ANT
75
| | ET50 | U80-20 |
| --- | --- | --- |
| Mock | 14.05191 | 2.5027 |
| 10uM ANT | 14.90137 | 2.94847 |
% Germination
50
25
0
7
10
20
Time (hrs)
Supplemental Figure 6 ANT does not alter ET_50_ or uniformity of L. sativa ‘Salinas’ seeds. Germination data was fit to the best fit time to event model in drc package. (A) Vertical dashed lines indicate ET_50_ values. (B) U_80-20_ is not affected by ANT application. ET_50_ values are normalized and plotted on the same point on the x axis. Vertical lines represent ET_20_ and ET_80_.

### Slide 7
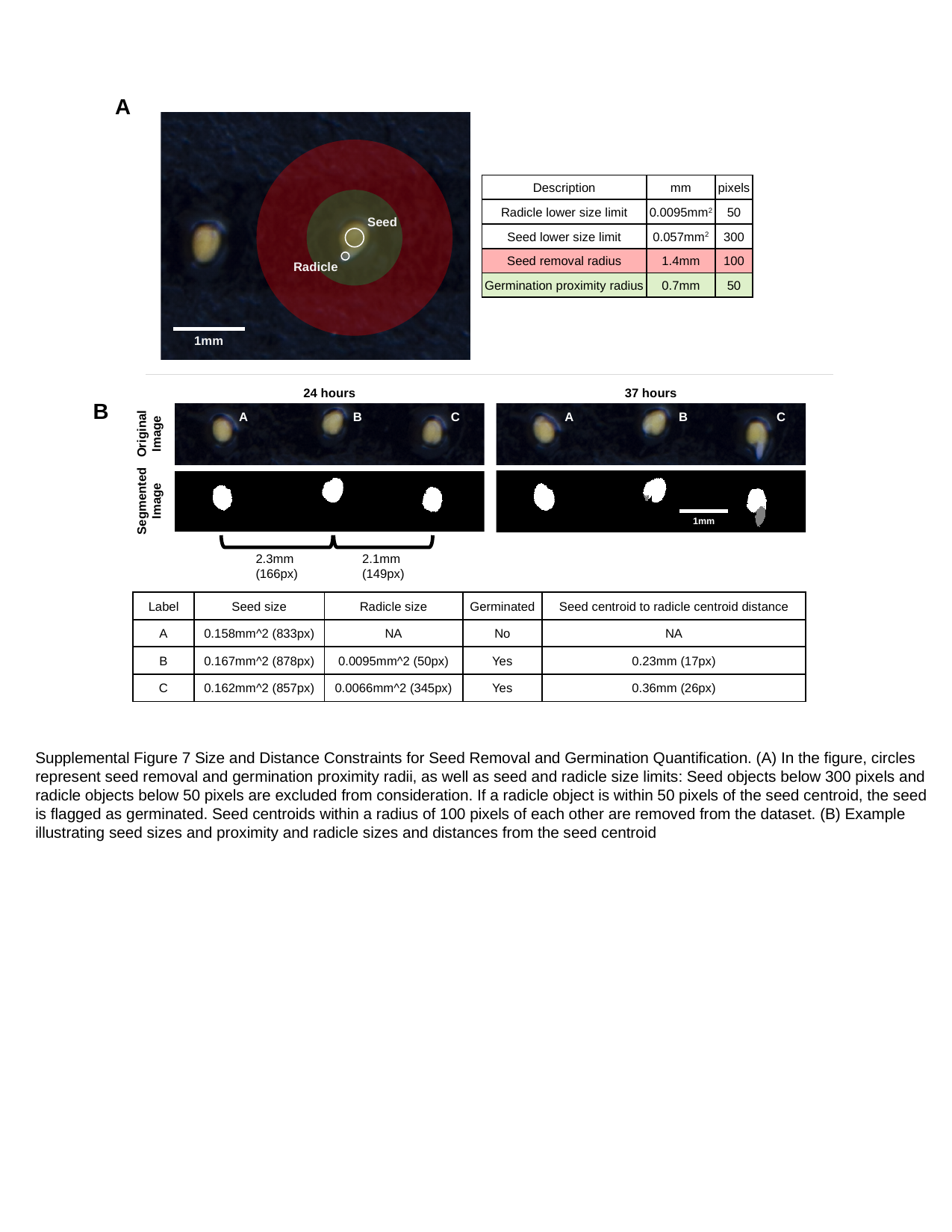

A
| Description | mm | pixels |
| --- | --- | --- |
| Radicle lower size limit | 0.0095mm^2^ | 50 |
| Seed lower size limit | 0.057mm^2^ | 300 |
| Seed removal radius | 1.4mm | 100 |
| Germination proximity radius | 0.7mm | 50 |
Seed
Radicle
1mm
24 hours
37 hours
B
A
B
C
A
B
C
Original Image
Segmented Image
1mm
2.3mm
(166px)
2.1mm
(149px)
| Label | Seed size | Radicle size | Germinated | Seed centroid to radicle centroid distance |
| --- | --- | --- | --- | --- |
| A | 0.158mm^2 (833px) | NA | No | NA |
| B | 0.167mm^2 (878px) | 0.0095mm^2 (50px) | Yes | 0.23mm (17px) |
| C | 0.162mm^2 (857px) | 0.0066mm^2 (345px) | Yes | 0.36mm (26px) |
Supplemental Figure 7 Size and Distance Constraints for Seed Removal and Germination Quantification. (A) In the figure, circles represent seed removal and germination proximity radii, as well as seed and radicle size limits: Seed objects below 300 pixels and radicle objects below 50 pixels are excluded from consideration. If a radicle object is within 50 pixels of the seed centroid, the seed is flagged as germinated. Seed centroids within a radius of 100 pixels of each other are removed from the dataset. (B) Example illustrating seed sizes and proximity and radicle sizes and distances from the seed centroid
